# Topical application of the cold-mimetic L-menthol decreases wheel running without affecting the beneficial effects of voluntary exercise in mice

**DOI:** 10.1101/2025.02.21.639574

**Authors:** Annalaura Bellucci, Bradley J. Baranowski, Stewart Jeromson, Michael Akcan, Serena Trang, Meagan Arbeau, Hadil Alfares, Katelyn Eisner, David C. Wright

## Abstract

Topical application of L-menthol, a pharmacological cold-mimetic and agonist of the cold-sensing receptor TRPM8 (Transient Receptor Potential Cation Channel Subfamily M Member 8), has been shown to stimulate brown adipose tissue (BAT) thermogenesis and reduce weight gain in both obese and lean male mice, without affecting energy intake. While these findings suggest that L-menthol could offer a novel approach to prevent weight gain, its potential to enhance the benefits of exercise on whole-body metabolic health remains unexplored. In this study, we investigated whether daily topical L-menthol application, combined with voluntary wheel running, could enhance exercise-induced improvements in metabolic health in male and female C57BL/6J mice housed at thermoneutrality (29°C). Our results demonstrated that although L-menthol treatment reduced voluntary wheel running distance there was still a main effect of exercise to reduce fat mass, weight gain and improve glucose tolerance. Indirect calorimetry revealed that L-menthol increased total energy expenditure, potentially explaining improvements in metabolic health despite reductions in voluntary wheel running. These findings suggest that although L-menthol does not enhance the effects of voluntary exercise, it remains a promising strategy for improving metabolic health.

**Key points:**

- L-menthol treatments led to a reduction in voluntary wheel running distance
- Despite the reduced voluntary exercise with L-menthol, wheel running led to significant reductions in fat mass as well as improved glucose tolerance
- Treatment with L-menthol increased energy expenditure perhaps providing an explanation for exercise-induced improvements in indices of metabolic health despite reduction in wheel running

## Introduction

Transient receptor potential (TRP) channels, a family of thermoreceptors, are primarily expressed in afferent dorsal root ganglion (DRG) sensory neurons(1) and trigeminal ganglia that innervate the skin, functioning to relay environmental temperature changes to the central nervous system (CNS)(2). Among these channels, the transient receptor potential cation channel subfamily M member 8 (TRPM8) plays a key role in cold-sensing through the activation of brown adipose tissue (BAT) thermogenesis (3). TRPM8 is activated by innocuous cooling temperatures below 30°C and is essential for detecting unpleasant cold stimuli or mediating the effects of cold-induced analgesia(4). Additionally, it serves as a receptor for chemical ligands, such as L-menthol and icilin, which are naturally derived and synthetic cooling compounds, respectively(5). In C57BL/6J male mice housed at thermoneutrality, activation of TRPM8 via topical L-menthol application has been shown to stimulate BAT thermogenesis, attenuate weight gain, and increase energy expenditure through a UCP1 and norepinephrine-dependent mechanism(6). In human studies a single topical application of L-menthol has been shown to increase thermogenesis(7), metabolic rate, and exercise performance, particularly in hot environments(8–11). In young, healthy men, acute oral administration of L-menthol has been shown to induce hyperventilation, improve endurance capacity, and lower the rate of perceived exertion (RPE)(12). Similarly, in well-trained male runners, L-menthol ingestion improved breathing comfort and boosted endurance capacity during exhaustive endurance running(13). Furthermore, a meta-analysis of randomized control trials found that L-menthol promoted a cooler thermal sensation and improved thermal comfort during exercise(14).

Factors such as environmental conditions, body mass index, and route of application have been identified as key contributors to enhancing the effects of L-menthol on endurance performance in humans(14). Given the promising evidence that topical application of the pharmacological cold-mimetic L-menthol mitigates weight gain in rodents and enhances thermogenesis, metabolic rate and exercise performance in humans, it is compelling to explore whether topical L-menthol application can potentiate the metabolic benefits of exercise. Therefore, the aim of this study was to investigate the impact of topical L-menthol treatment on the effects of voluntary physical activity in mice. We hypothesized that menthol treatment would potentiate the metabolic effects of voluntary wheel running in mice of both sexes housed at thermoneutrality.

## Methods

### Animals and ethics

All protocols were approved by the University of British Columbia Animal Care Committee (protocol# A22-0011) and followed Canadian Council on Animal Care Guidelines. Male and female C57BL/6J mice (∼12-14 weeks of age) (CAT# 000664, Jackson Laboratories) were group housed (∼4/cage) at room temperature (20-22°C) and given ad libitum access to standard chow (CAT# 2918, Teklad) and water while acclimating to the animal facility at BC Children’s Hospital Research Institute for 1 week. Thereafter mice were single housed in shoe-box cages, given ad libitum access to standard chow diet and water and acclimated at thermoneutrality (29°C) in the Solace Zone Caging System (Alternative Design Manufacturing & Supply Inc.).

### Topical L-menthol treatments and voluntary wheel running

After 1 week acclimation at thermoneutrality (29°C), 12-14 weeks-old C57BL/6J female and male mice were weight-matched, and given access to a running wheel or remained sedentary for 3 weeks and treated daily with topical L-menthol (2g/kg BW, 5% wt/vol CAT# W266590 Sigma– Aldrich) or ethanol control(6). L-menthol or ethanol were selectively applied to the unshaved dorsal surface of restrained mice carefully avoiding orifices, limbs, and the tail as previously described(6). All treatments occurred between 10.00 am and 12.00 pm. Running distance was recorded daily, and body weight and food intake were measured weekly.

Pre and post intervention, body composition was measured using an EchoMRI body composition analyzer (Echo MRI, Huston TX). Tissue collections occurred 24 hours after the last L-menthol / ethanol control treatments. Running wheels were locked the night before (∼8 pm) and mice euthanized after a 6hrs fast starting at ∼07.00 am. In the second set of experiments 12-14 weeks-old C57BL/6J female mice housed at thermoneutrality (29°C) were given access to a running wheel or remained sedentary for 2 weeks and treated daily with a relative dose (2g/Kg) of topical L-menthol (5% wt/vol) or the ehtanol control. On day 15 the last L-menthol / ethanol control treatments occurred between 10.00 am and 12.00 pm and mice were euthanized and tissue harvested after 6 hours of voluntary wheel running activity in the dark cycle following their peak running activity (∼12 am).

### Oral glucose tolerance test

Glucose tolerance was assessed ∼ 24 hours after the last topical L-menthol / ethanol control treatment during the third week of intervention, and running wheels were locked at 6 pm the day before the OGTT. Mice were fasted for 6 h (starting at ∼07.00 am) and given an oral gavage of glucose (2 g/Kg BW)(15). Blood glucose was taken from a small drop of blood from the tail vein using a handheld glucometer and glucose strips (Freestyle Lite; Abbott Laboratories, Abbott Park, Illinois, USA) and measured immediately pre-gavage (time point 0) as well as 15-, 30-, 60-, 90- and 120-min post-gavage.

### Acute topical L-menthol treatment and forced treadmill running

12-14 weeks-old C57BL/6J female mice housed at thermoneutrality (29°C) were weight-matched and assigned to either L-menthol / ethanol control and exercised / sedentary groups. Mice in the exercise groups were familiarized to a motorized rodent treadmill (Exer 3/6 Columbus Instruments) for two consecutive days for 15 min / day at 15 m / min on a 5% incline and then given a rest period of ∼48 hours before maximum running speed was assessed as previously described(16,17). Maximum speed was determined by running mice at 10 m / min for 3 minutes at a 5% incline and the speed was subsequently increased by 3 m / min every 3 minutes. Maximum speed was defined as the fastest speed mice could maintain for 3 continuous minutes.

Approximately 48 hours following the maximum speed test, mice received an acute topical L-menthol or ethanol control treatment between 10.00 am and 12.00 pm.

The exercise groups ran for 2 hours in the dark cycle (start ∼6pm), or until exhaustion, at 70% of their predetermined maximum speed with an incline of 5%. The sedentary mice were in the same procedure room as the exercised mice when conducting treadmill acclimation, maximum speed test and 2 hours of acute treadmill running.

### Indirect calorimetry measurements

Prior to the beginning of the metabolic cage measurements, 12-14-week-old C57BL/6J female mice, housed at thermoneutrality (29°C), were treated daily with either a topical application of L-menthol (2 g/kg) or an ethanol control. After a 2-week treatment period, mice were individually housed in metabolic cages (Promethion High-Definition Multiplexed Respirometry System for Mice; Sable Systems International, Las Vegas, NV, USA) at thermoneutrality (29°C), with ad libitum access to food and water. Mice were allowed to acclimate for 24 hours before collecting data over a full light and dark cycle. Mice continued receiving their respective daily treatments of either L-menthol or ethanol throughout the duration. Respirometry values were recorded every 5 minutes, with a 30-second dwell time per cage and a baseline cage sampling frequency of 30 seconds occurring every four cages. The respiratory exchange ratio (RER) was calculated as the ratio of VCO_2_ to VO_2_. Locomotion was measured based on beam breaks within a grid of infrared sensors in each cage. Energy expenditure was calculated using the Weir equation(18): Energy expenditure = 3.941 kcal/L × VO_2_ + 1.106 kcal/L × VCO_2_. After 48 hours, mice were euthanized, and tissues harvested.

### Tissue collection

Mice were anesthetized with pentobarbital (Euthansol 120 mg / Kg BW, IP) and euthanized through exsanguination(19). Blood was collected via cardiac puncture, immediately placed on ice for ∼30 min prior to centrifugation (1500 x g for 10 minutes) and serum was collected. The liver, inguinal white adipose (iWAT), epididymal white adipose tissue (eWAT), gonadal white adipose tissue (gWAT), BAT, hypothalamus, and triceps muscles were harvested, snap frozen in liquid nitrogen and stored at -80° C until further analysis.

### Serum Analysis

Metabolites and hormone assays were analyzed with the Versamax Tunable Microplate Reader and SoftMax Pro Software (Molecular Devices). Insulin (CAT# 10-1247-01, Mercodia Inc.) and corticosterone (CAT# 55-CORMS-E01, Alpco, Salem) were measured with commercially available enzyme-linked immunosorbent assay (ELISA) kits. Serum glucose (CAT# 10009582, Cayman Chemicals), NEFA (Non -Esterified Fatty Acids) (CAT# CA97000-012, Wako Chemicals) and BHB (Beta-Hydroxybutyrate) (CAT# 700190, Cayman Chemicals) were measured using commercially available colorimetric assay kits.

### Real-time PCR

RNA was extracted from the hypothalamus using Trizol and Qiagen RNeasy Mini Kits (CAT# 74106) followed by DNase-free treatment (Thermo Fisher Scientific, CAT# AM1906) for the removal of genomic DNA. cDNA was synthesized using Superscript II (CAT# 4368814, Thermo Fisher Scientific) and real-time PCR was run using SYBR Green Supermix (CAT# 1725271, Biorad) on a Bio-Rad CFX connect system. Gene expression was measured relative to the housekeeping genes, β-actin *(Actb)* and *Ppib* with the 2^−ΔΔ^CT method as previously described. β-actin (*Actb*) and *Ppib* have been shown to be suitable housekeeping genes (20,21) and in the current study we found that raw CT values were not different between groups. Primer sequences were as follows: *Actb:* Rev 5’-GAGCATAGGCCTCGTAGAT-3’, Fwd 5’-GACCCAGATCATGTTTGAGA-3’ ; *Ppib:* Rev 5’-GCCCGTAGTGCTTCAGCTT-3’, Fwd 5’-GGAGATGGCACAGGAGGAA-3’ ; *AgRP*: Rev 5’-GGTACCTGCTGTCCAAAGCAG-3’, Fwd 5’-AGTCTGACTGCATGTTGCGT-3’ ; *Orexin*: Rev 5’-GTTCGTAGAGACGGCAGGAA-3’, Fwd 5’-ACTTTCCTTCTACTACAAAGGTTCCCT-3’; *Npy*: Rev 5’-TGTCGCAGAGCGGAGTAGTAT-3’, Fwd 5’-ATGCTAGGTAACAAGCGAATGG-3’

### Liver glycogen

Glycogen concentrations were measured in liver as described by Schaubroeck et al(22). Briefly, tissue was homogenized in 0.5 M NaOH followed by heating at 100°C for 30 minutes with periodic mixing. Na_2_SO_4_ and ethanol were added to the tissue homogenate, samples were centrifuged at 2,000 x g for 10 minutes to precipitate glycogen and then resuspended in ddH2O.

Sulphuric acid and phenol were added to 50ul of sample and the reaction proceeded for 30 minutes. Samples were then transferred to a 96-well plate and absorbance was read at 488 nM.

### Statistical Analysis

Statistical tests were completed using GraphPad Prism v.10.0 (GraphPad Software, La Jolla, CA, USA). Body weight gain, body composition, total food intake, glucose area under the curve and tissue weights were analyzed by two-way ANOVA. Tukey’s post-hoc analysis was conducted if there was a significant interaction between L-menthol treatment and voluntary wheel running/treadmill running. Wheel and treadmill running distances as well as the time spent running until exhaustion, energy expenditure, oxygen consumption, and carbon dioxide emission, were analyzed using unpaired t-tests, two-tailed T-tests, or non-parametric t-tests when data was not normally distributed. Pearson correlation coefficient analysis was used to examine associations between serum corticosterone levels and running distance. HOMA-IR (Homeostatic Model Assessment of Insulin Resistance) was calculated based on fasting blood glucose and fasting insulin levels as reported (23). Data are presented as means ± SD, and individual data points are shown when possible. A relationship was considered significant when p < 0.05.

## Results

### 3 weeks of topical L-menthol treatment reduce voluntary wheel running distance in female and male mice

We were interested in examining the combined effects of menthol treatment and wheel running on weight gain and indices of glucose metabolism in mice. At first, we assessed the effects of menthol treatment on wheel running performance. As shown in Figure 1 daily L-menthol treatment significantly decreased voluntary wheel running distance in both female (Figure 1A and 1B) (p=0.0161) and male mice (p=0.0451) (Figure 1C and 1D). Additionaly, the gene expression of neuropeptides linked to wheel running behaviour (24–27) including *Orexin* (exercise: p=0.7537; treatment: p=0.3505; interaction: p=0.2393), *Npy* (exercise:p=0.0864; treatment: p=0.4943; interaction: p=0.6262) and *AgRP* (exercise: p=0.7454; treatment: p=0.7454; interaction: p=0.1397) in the hypothalamus were not different between groups (Table 1).

**Figure 1:**
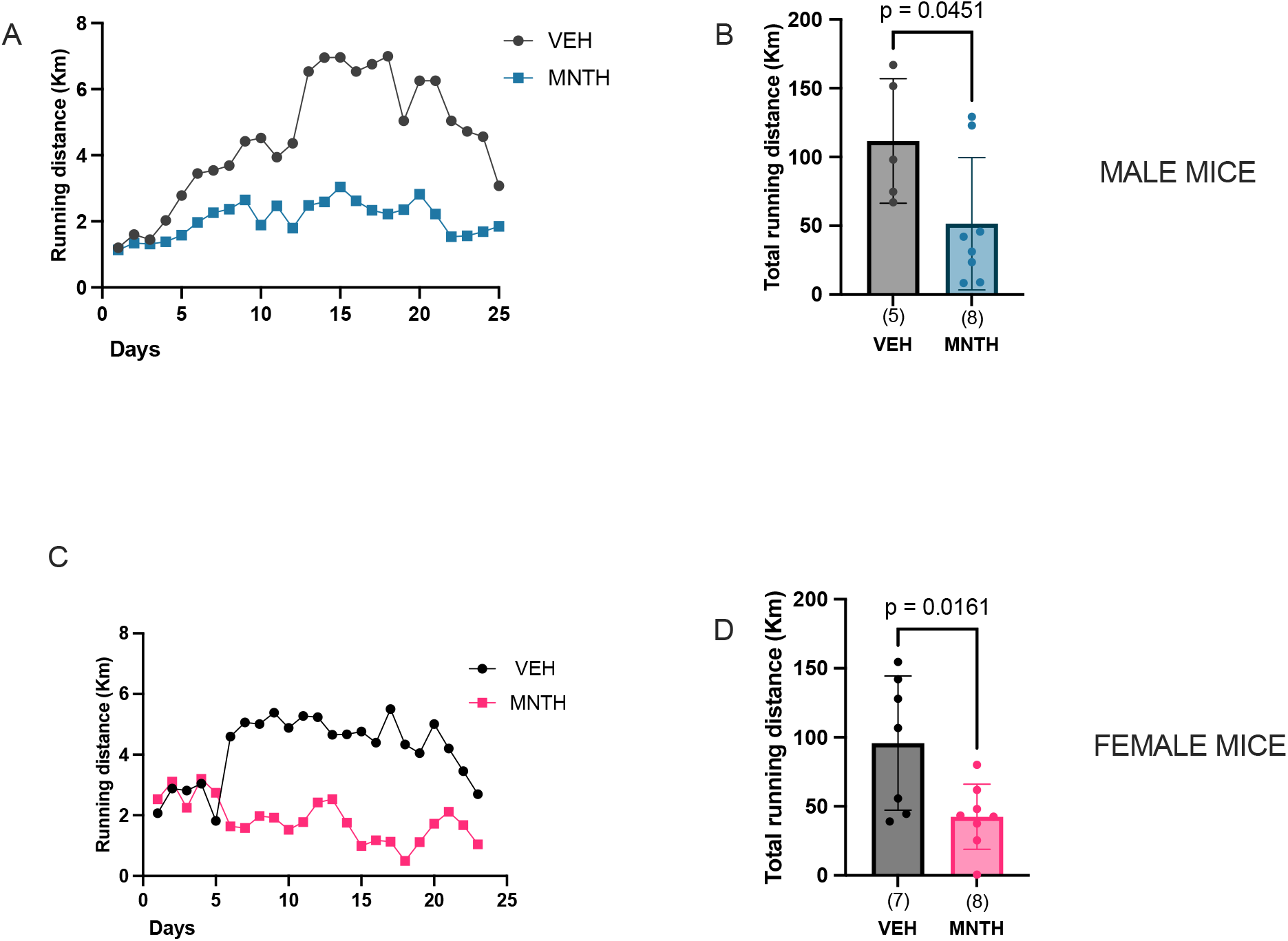
3 weeks of topical L-menthol treatment reduces voluntary wheel running distance in female and male mice. 3 weeks of topical L-menthol treatments decreased voluntary wheel running distance in female (B) and male mice (D). (A) daily running distance of female mice and (B) total running distance; (C) daily running distance of male mice and (D) total running distance. Data are presented as means ± SD in C and D with individual data points shown. Numbers per group are shown in parenthesis below each bar. Data were analyzed using unpaired, two tailed T-tests in B and a Mann-Whtiney test in D. Bars joined by lines are significant at the p-value that is shown.

**Table 1:** L-menthol does not alter hypothalamic orexigenic neuropeptides gene expression. mRNA expression of *Orexin, Npy* and *AgRP* were analayzed in the hypothalmus of female mice after 3 weeks of voluntary wheel running and L-menthol treatment. Two way-ANOVA (exercise x treatment) was used to determine differences between groups. Numbers of mice per group are shown in parentesis. Data are presented as mean ± SD.

|  | SED VEH | EX VEH | SED MNTH | EX MNTH | p-value |
| --- | --- | --- | --- | --- | --- |
| ***Orexin*** | 1.000±0.298(8) | 1.141±0.266(7) | 1.200±0.25(8) | 1.117±0.209(8) | Exercise:0.7537<br>Treatment:0.3505<br>Interaction:0.2393 |
| ***Npy*** | 1.000±0.235(8) | 1.106±0.246(7) | 1.016±0.236(8) | 1.204±0.200(8) | Exercise:0.0864<br>Treatment:0.4943<br>Interaction:0.6262 |
| ***AgRP*** | 1.000±0.517(8) | 1.193±0.533(7) | 1.300±0.424(8) | 1.000±0.302(8) | Exercise:0.0864<br>Treatment:0.4943<br>Interaction:0.6262 |

### 3 weeks of voluntary exercise, but not L-menthol treatment, improves indices of glucose homeostasis

We next aimed to investigate whether L-menthol would enhance the beneficial effects of exercise on indices of glucose metabolism. We found that daily voluntary wheel running, but not L-menthol treatment, improved glucose tolerance in both female (Figure 2A and 2B) (exercise: p=0.0199; treatment: p=0.6166; interaction:p=0.7611) and male mice (Figure 2F and 2G) (exercise: p=0.0051; treatment: p = 0.1599; interaction: p=0.4523). Fasting blood glucose was reduced in the exercised female (exercise: p=0.0046; treatment: p=0.9082; interaction: p=0.7283) (Figure 2C) and male mice (exercise:p=0.0001; treatment: p=0.4822; interaction: p=0.0782) (Figure 2H). Serum insulin levels were not reduced in either female (Figure 2D) (exercise: p=0.0914; treatment: p=0.613; interaction: p=0.9413) or male mice (Figure 2I) (exercise: p=0.5812; treatment: p=0.3767; interaction p=0.7562) with exercise. The homeostasis model assessment-estimated insulin resistance (HOMA-IR) was reduced in both female (Figure 2E) (exercise p=0.0355; treatment p=0.8086; interaction: p=0.8854) and male (Figure 2L) (exercise p=0.0394; treatment: 0.3352; interaction: p=0.6682) mice with exercise.

**Figure 2:**
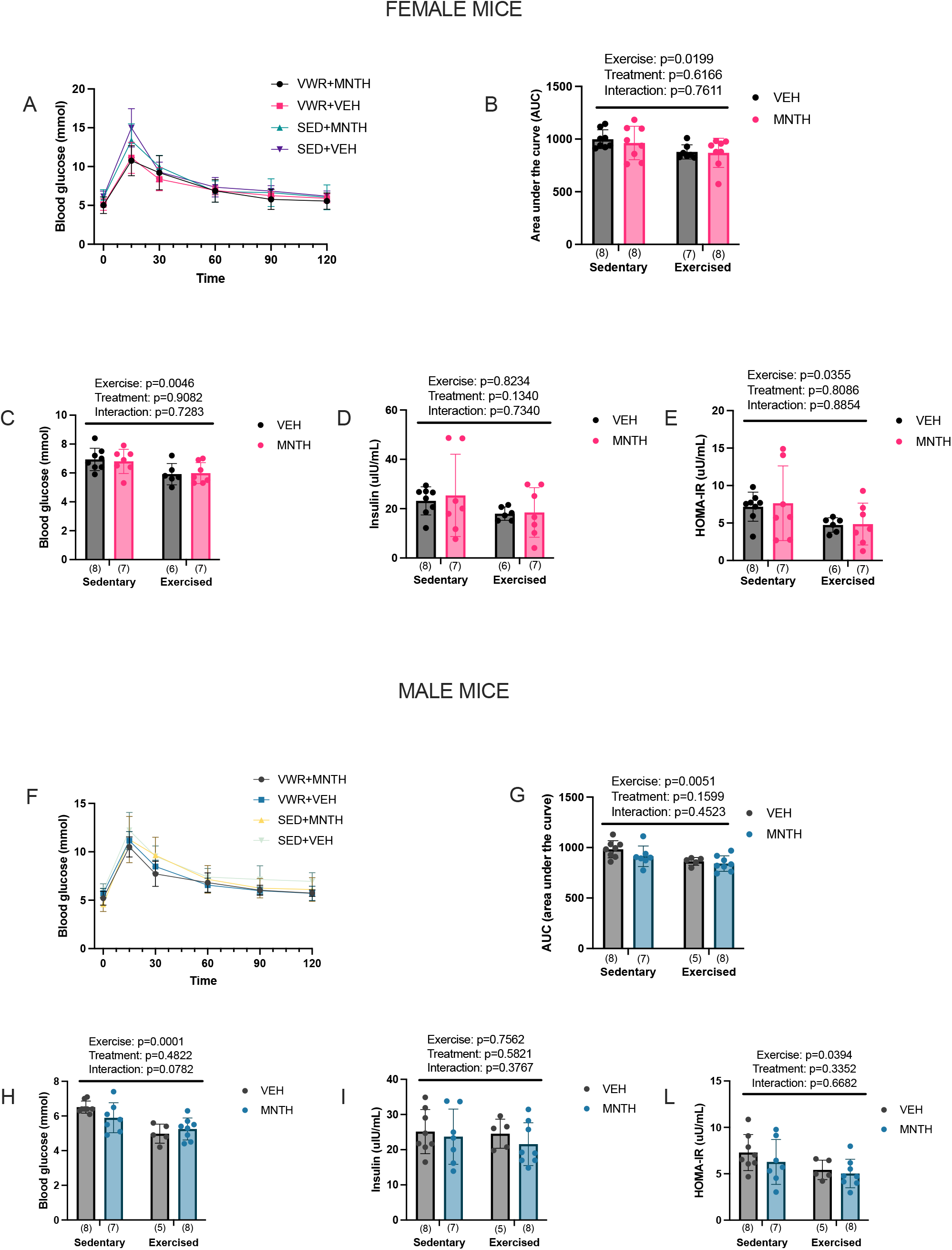
3 weeks of voluntary exercise, but not L-menthol treatment, improves indices of glucose homeostasis. 3 weeks of voluntary wheel running improves glucose tolerance (B), lowers fasting blood glucose (C,H) and HOMA-IR (E,L) in female and male mice. (A) and (F) show glucose curves over time during the glucose tolerance test. Two way-ANOVA (exercise x treatment) was used to determine differences between groups with main and interaction effects shown as an inset. Data are presented as mean ± SD with individual data points shown. Numbers per group are shown in parenthesis below each bar.

### Exercise reduces fat mass in female and male mice

It has previously shown that topical L-menthol treatments reduce body weight gain in both obese and lean mice housed at thermoneutrality(6). In this experiment, exercise did not attenuate weight gain in female mice (Figure 3A) (exercise: p=0.1363; treatment: p=0.7240; interaction: p=0.1363), whereas it attenuated weight gain in male mice (Figure 3G) (exercise: p=0.0104; treatment: p=0.2207; interaction: p=0.6832).

**Figure 3:**
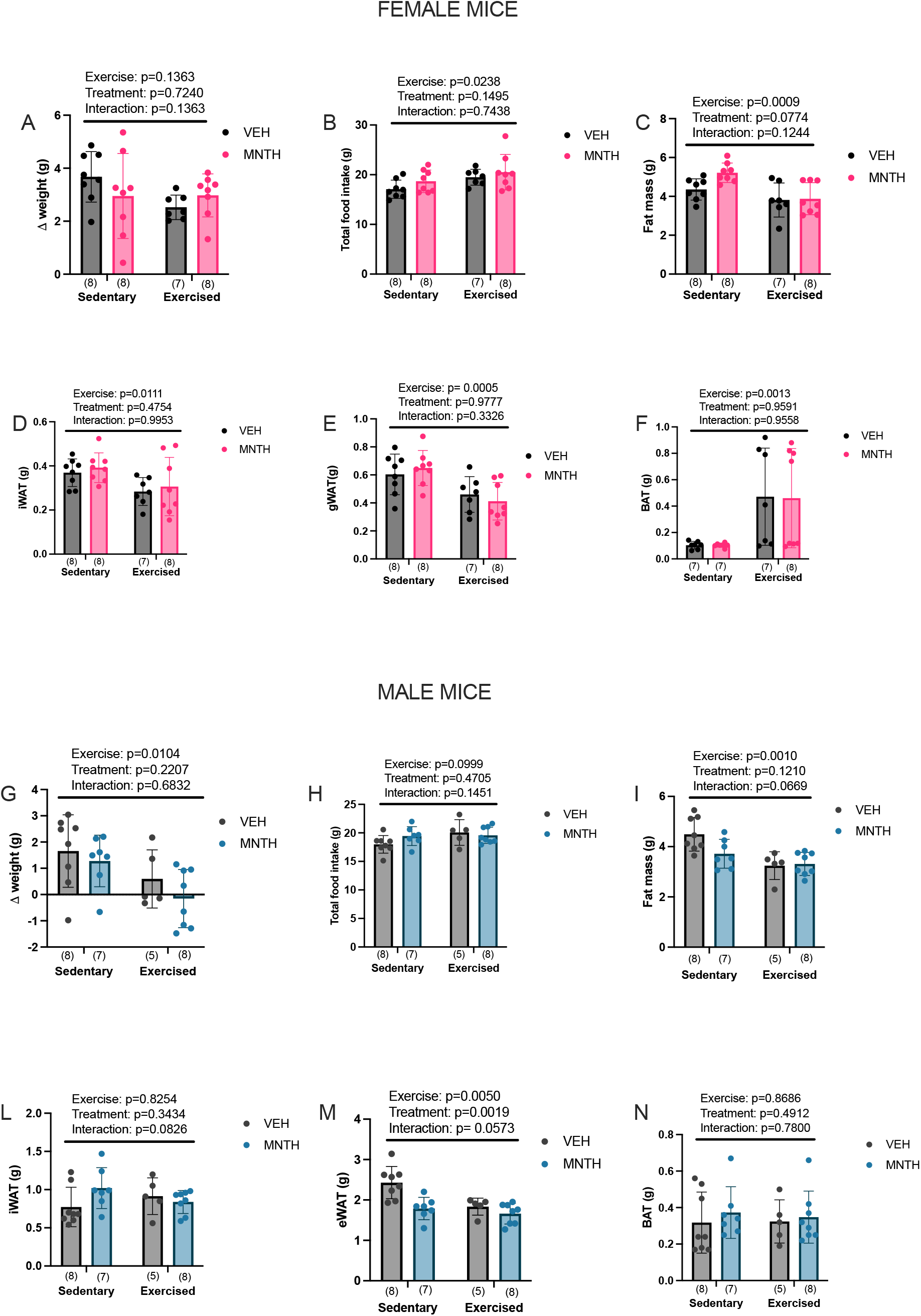
Exercise reduces fat mass in female and male mice. Exercise does not attenuate weight gain (A) but increases food intake in female mice (B), while attenuating weight gain (G) independent of food intake (H) in male mice. Exercise decreases fat mass in female (C) and male mice (I) and reduces iWAT (D) and gWAT (E) mass in female mice. Menthol and exercise reduce eWAT (M) but not iWAT mass (L) in male mice, while exercise increases BAT mass in female (F) but not in male mice (N). Two way-ANOVA (exercise x treatment) was used to determine differences between groups with main and interaction effects shown as an inset. Data are presented as mean ± SD with individual data points shown. Numbers per group are shown in parenthesis below each bar.

Exercise increased total food intake in female (Figure 3B) (exercise: p=0.0238; treatment: p=0.1495; interaction: p=0.7438) but not male mice (Figure 3H) (exercise: p=0.0999; treatment: 0.4750; interaction: p=0.1451). Exercise reduced fat mass in female mice (Figure 3C) (exercise: p=0.0009; treatment: p=0.0774; interaction: p=0.1244) and male mice (Figure 3I) (exercise: p=0.0010; interaction: p=0.0669; treatment: p=0.1210). L-menthol treatment and exercise reduced eWAT mass (Figure 3M) (exercise p=0.0050; treatment: p=0.0019; interaction: p=0.0573) but not iWAT mass in male mice (Figure 3L) (exercise: p=0.8254; treatment: p=0.3434; interaction: p=0.0826), whereas in female mice exercise significantly reduced iWAT (Figure 3D) (exercise: p=0.0111; treatment: p=0.4754; interaction: p=0.9953) and gWAT mass (Figure 3E) (exercise: p=0.0005; treatment: p=0.9777; interaction: p=0.3326). Finally, exercise increased BAT mass in female mice (Figure 3F) (exercise: p=0.0013; treatment: p=0.9591; interaction: p=0.9558) but not in male mice (Figure 3N) (exercise: p=0.8686; treatment: p=0.4912; interaction: p=0.7800).

### Topical application of L-menthol increase energy expenditure

A previous study demonstrated that acute topical application of L-menthol increases oxygen consumption, energy expenditure, and core body temperature in male mice, through a TRPM8 dependent mechanism(6). We found that L-menthol treatment increased total energy expenditure (Figure 4A) (p = 0.0021), oxygen consumption (Figure 4B) (p = 0.0018), and carbon dioxide production (Figure 4C) (p = 0.0054), whereas RER (Figure 4D) (p = 0.9644) and total locomotion (Figure 4E) p = 0.3987) were not affected by L-menthol treatment.

**Figure 4:**
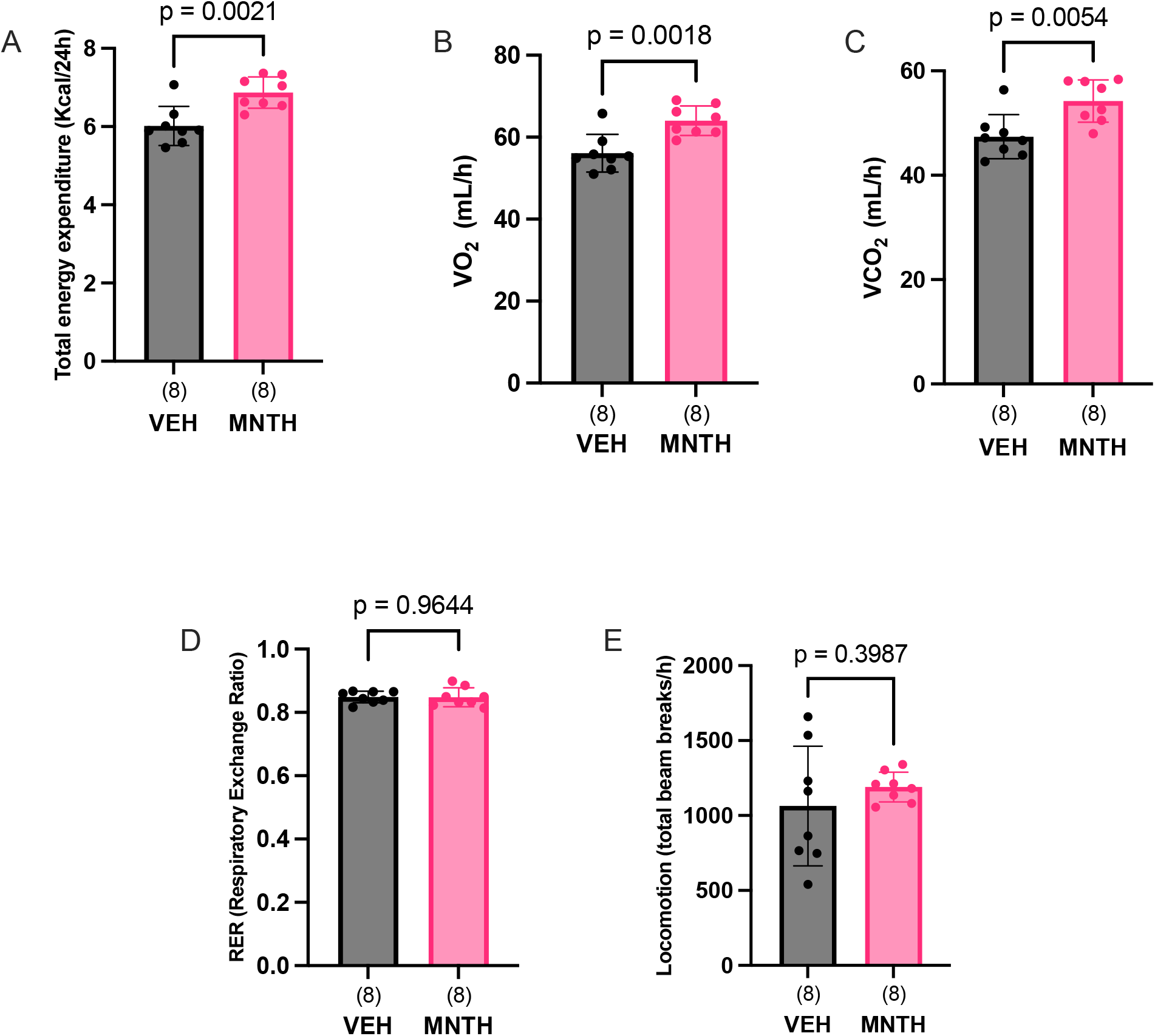
Topical application of L-menthol increase energy expenditure. Total energy expenditure (A), oxygen consumption (VO_2_) (B), and carbon dioxide production (VCO_2_) (C) are increased in menthol-treated mice. RER (Respiratory Exchange Ratio) (D) and total locomotion (E) did not differ between vehicle and menthol treated-mice (D). Data were analyzed using unpaired, two tailed T-tests. Data are presented as mean ± SD with individual data points shown. Numbers per group are shown in parenthesis below each bar.

### Indices of glucose homeostasis are improved following peak wheel running activity and L-menthol increases serum corticosterone

In our initial experiments, endpoints were measured approximately 24 hours after the last L-menthol treatment, with running wheels locked the evening prior. As a next step, we aimed to investigate whether L-menthol treatment influenced indices of glucose homeostasis when measured immediately following peak wheel running activity. We also sought to determine whether reductions in wheel running due to L-menthol treatment were reflected in differences in the hypothalamic gene expression of neuropeptides following peak wheel running activity. To do so, we repeated the menthol/wheel running experiment and harvested tissues following mice peak wheel running activity in the dark cycle (∼12:00 am). These experiments were only completed in female mice as they are better runners than males. We confirmed that topical L-menthol treatments reduced voluntary wheel running distance (Figure 5A and 5B) (p=0.0006) and that exercise significantly reduced weight gain (Figure 5C) (exercise: p = 0.0121; treatment: p=0.2680; interaction: p=0.8910), fat mass (Figure 5D) (exercise: p<0.0001; treatment:p=0.7866; interaction: p=0.1605), iWAT mass (Figure 5E) (exercise:p=0.0005; treatment:p=0.6114; interaction:p=0.1698) and gWAT mass (Figure 5F) (exercise p=0.0004; treatment:p=0.2037; interaction:p=0.1948). When measured immediately following peak wheel running activity, exercise decreased insulin levels (exercise: p=0.0151; treatment:p=0.4189; interaction: p=0.1944) (Figure 4H) and HOMA-IR (exercise: p=0.0116; treatment: p=0.6478; interaction: p=0.2744) (Figure 4I) while glucose levels did not differ between groups (exercise: p=0.8398; treatment: p=0.0780; interaction: p=0.3310) (Figure 4G). Furthermore, following peak wheel running activity, we observed increased *Orexin* (exercise: p=0.0247; treatment: p=0.8944; interaction: p=0.5565), and *Npy* (exercise:p=0.0010; treatment: p=0.2777; interaction: p=0.5276) but not *AgRP* (exercise: p=0.0870; treatment: p=0.7175; interaction: p=0.2241) mRNA expression (Table 2). Corticosterone is a stress hormone that increases in proportion to exercise intensity(28,29) and has previously been shown to reduce voluntary wheel running activity (30). We found that 2 weeks of topical L-menthol treatment increased serum corticosterone levels (Figure 5L) (exercise: p=0.4310; treatment: p=0.0154; interaction: p=0.9875), however serum corticosterone levels did not correlate with the distance ran on the wheel (Figure 5M) (r= 0.01015; p= 0.9725).

**Table 2:**
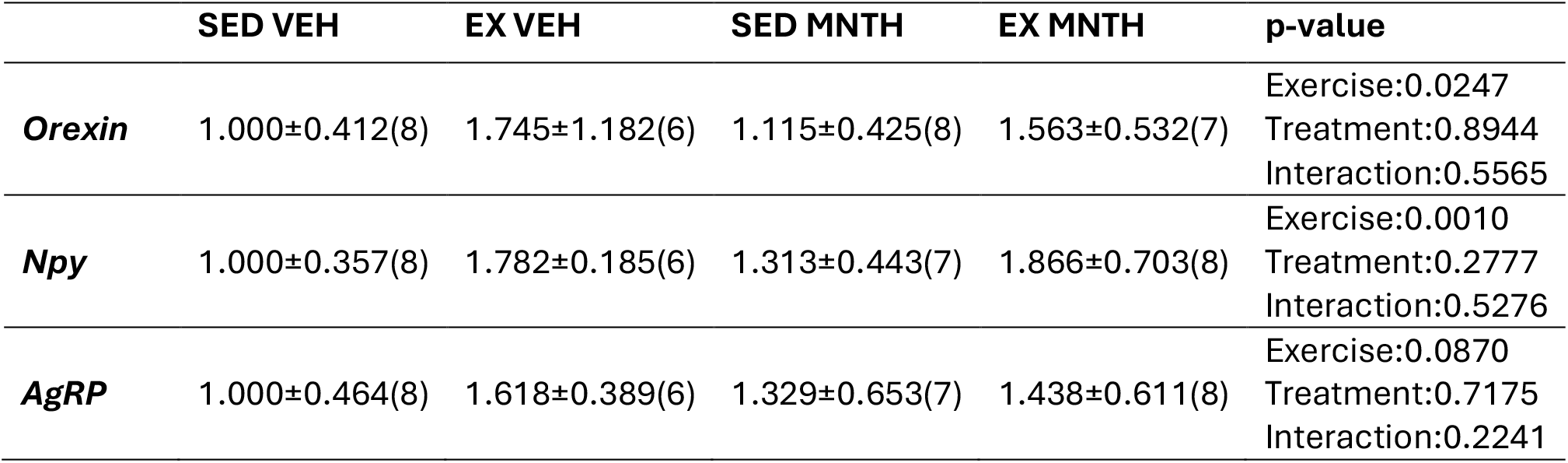
Hypothalamic orexigenic neuropeptides gene expression increases following peak wheel running activity. mRNA expression of *Orexin, Npy* and *AgRP* were analyzed in the the hypothalamus of female mice after peak wheel running activity. Two way-ANOVA (exercise x treatment) was used to determine differences between groups. Numbers of mice per group are shown in parentesis. Data are presented as mean ± SD.

|  | SED VEH | EX VEH | SED MNTH | EX MNTH | p-value |
| --- | --- | --- | --- | --- | --- |
| ***Orexin*** | 1.000±0.412(8) | 1.745±1.182(6) | 1.115±0.425(8) | 1.563±0.532(7) | Exercise:0.0247<br>Treatment:0.8944<br>Interaction:0.5565 |
| ***Npy*** | 1.000±0.357(8) | 1.782±0.185(6) | 1.313±0.443(7) | 1.866±0.703(8) | Exercise:0.0010<br>Treatment:0.2777<br>Interaction:0.5276 |
| ***AgRP*** | 1.000±0.464(8) | 1.618±0.389(6) | 1.329±0.653(7) | 1.438±0.611(8) | Exercise:0.0870<br>Treatment:0.7175<br>Interaction:0.2241 |

**Figure 5:**
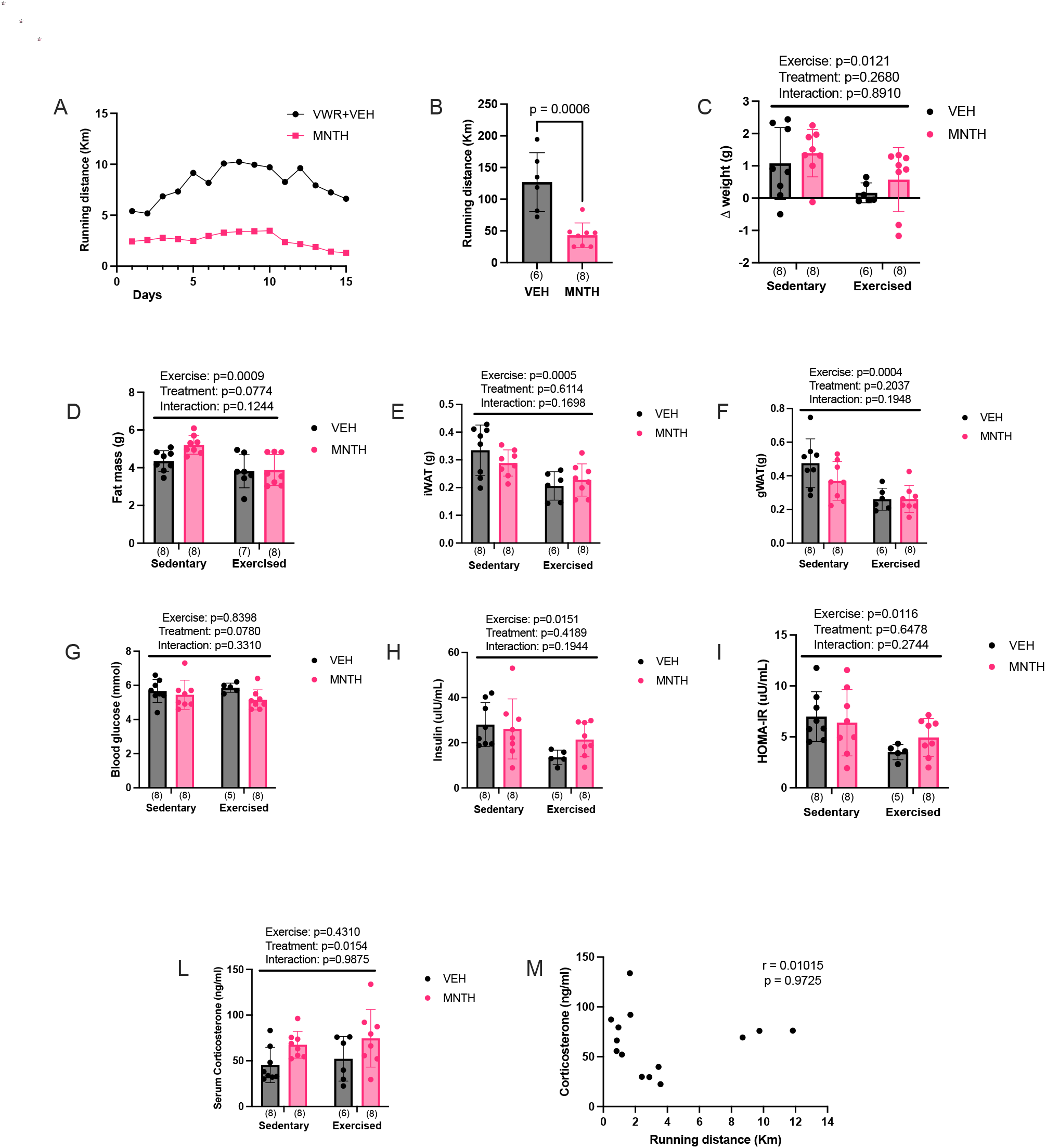
Indices of glucose homeostasis are improved following peak wheel running activity and L-menthol increases serum corticosterone. 2 weeks of L-menthol treatments reduced running distance (A,B) while exercise decreased weight gain (C), fat mass(D), iWAT(E) and gWAT(F). An acute bout of wheel running decreased insulin levels (H) and HOMA-IR (I). L-menthol increases serum corticosterone (L). Data were analyzed using unpaired, two tailed T-tests in B. Two way-ANOVA (exercise x treatment) was used to determine differences between groups with main and interaction effects shown as an inset. Data are presented as mean ± SD with individual data points shown. Numbers per group are shown in parenthesis below each bar.

### Topical L-menthol treatment does not attenuate distance run or the acute metabolic response to a bout of forced treadmill running

Our first experiments demonstrated that voluntary activity was reduced with L-menthol treatment. To investigate if this could be due to a physiological limitation we investigated if L-menthol reduces treadmill running to a similar extent as it does voluntary wheel running.

Female mice were treated with either L-menthol or ethanol control ∼ 8 hrs prior to an acute bout of treadmill exercise. We found that L-menthol had no effect on treadmill running distance (Figure 6A) (p=0.9791), nor did it alter the total time spent running compared to control mice (Figure 6B) (p=0.7428). The acute effects of exercise on circulating metabolites were generally the same between vehicle and menthol treated mice. Serum glucose levels were higher in the sedentary vehicle-treated group compared to the sedentary menthol-treated group (p=0.0277) (Figure 6C). Additionally, glucose levels were greater in the sedentary vehicle group compared to the exercised vehicle and menthol-treated groups (p<0.0001) (Figure 6C), and lower in the exercised vehicle-treated group compared to the sedentary menthol-treated group (p = 0.0028) (Figure 6C). Exercise increased circulating levels of NEFA (Figure6D)(exercise:p<0.0001; treatment:p=0.8171; interaction:p=0.5000), BHB (Figure6E)(exercise:p<0.0001; treatment:p=0.2597; interaction:p=0.5647) and corticosterone (Figure 6F) (exercise:p<0.0001; treatment:p=0.8142; interaction: p = 0.8262). Liver glycogen levels were significantly lower in the exercised menthol-treated group compared to the sedentary menthol-treated group (p<0.0001) (Figure 6G), and in the exercised vehicle-treated group compared to the sedentary vehicle-(p=0.0122) (Figure 6G) and menthol-treated groups (p<0.0001) (Figure 6G). We also observed a two-fold difference in liver glycogen levels between the sedentary menthol and sedentary vehicle-treated groups (p<0.0001) (Figure 6G). However, liver glycogen levels did not differ between the exercised menthol and exercised vehicle groups (p = 0.7500) (Figure 6G).

**Figure 6:**
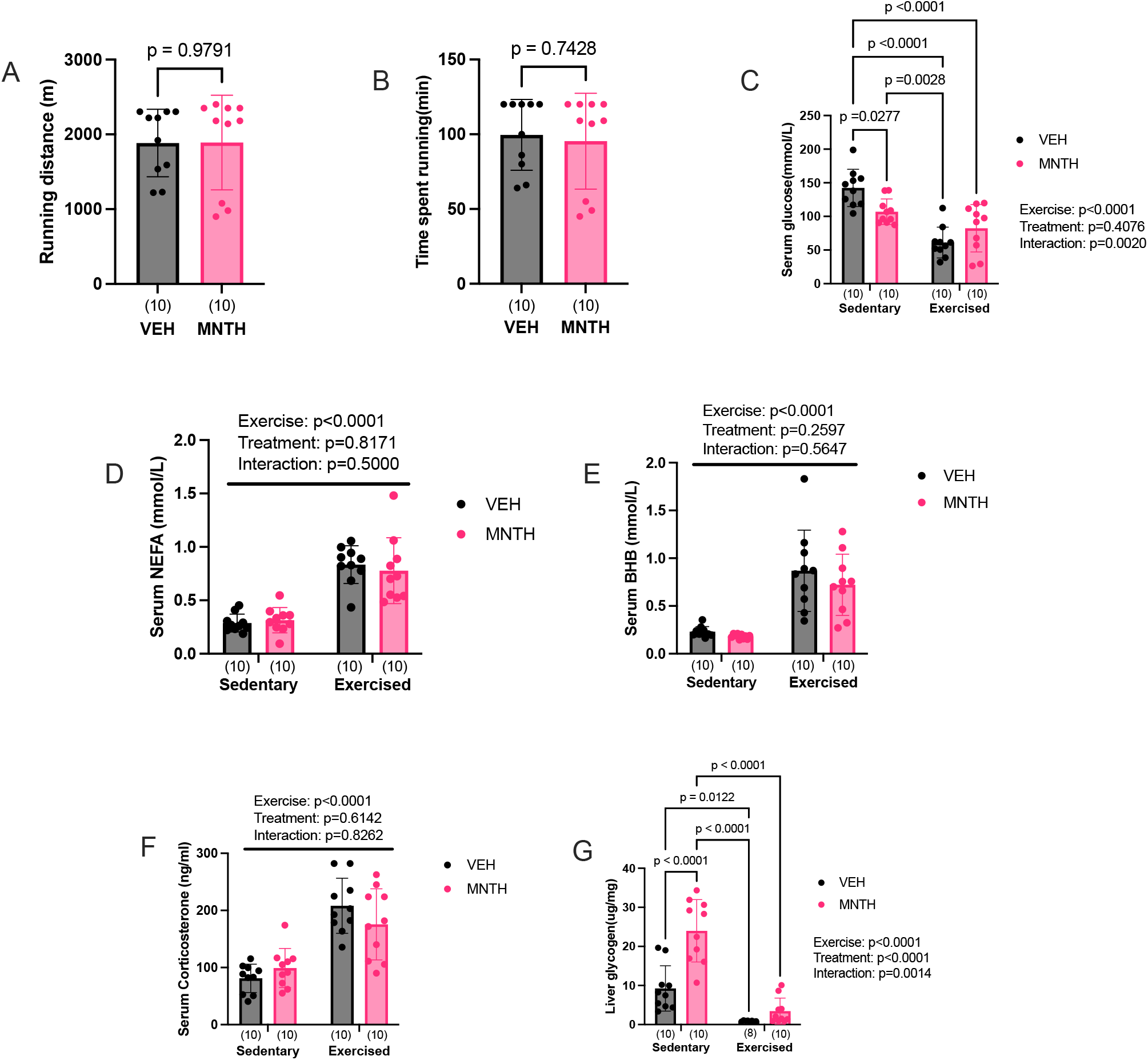
Topical L-menthol treatment does not attenuate distance run or the acute metabolic response to a bout of forced treadmill running. Treadmill running distance (A) and time to exhaustion (B) are not affected by an acute topical menthol treatment. Exercise reduces circulating levels of serum glucose (C) and increases serum NEFA(D), BHB(E), and corticosterone (F) levels. Exercise and menthol affected liver glycogen levels (G). Data were analyzed using unpaired, two tailed T-tests in A and B. Two way-ANOVA (exercise x treatment) was used to determine differences between groups with main and interaction effects shown as an inset. Tukey’s post-hoc test was used if a significant (exercise x treatment) interaction was found (C and G). Data are presented as mean ± SD with individual data points shown. Bars joined by lines are significantly different at the p-level shown. Numbers per group are shown in parenthesis below each bar.

## Discussion

The aim of this study was to determine whether topical application of the TRPM8 agonist and pharmacological cold-mimetic L-menthol, would enhance the beneficial effects of voluntary exercise in C57BL/6J mice housed at thermoneutrality. In the first set of experiments, using male and female mice we found that 3 weeks of L-menthol treatment reduced voluntary wheel running distance in both sexes. Despite the reduction in running activity with L-menthol, voluntary exercise improved glucose tolerance, reduced fat mass and blunted weight gain. Similar improvements in indices of glucose metabolism were observed, despite the reductions in wheel running caused by L-menthol treatment, when assessed immediately following the period of peak wheel running activity. Additionally, the decrease in wheel running with L-menthol treatment did not appear to be related to changes in hypothalamic neuropeptide gene expression or corticosterone levels (30,31). Since L-menthol reduced voluntary wheel running, we sought to determine whether this could be due to physiological limitations. To do so, we treated mice with menthol and assessed their treadmill running performance. In contrast to wheel running, there were no differences in running distance or time-to-exhaustion between L-menthol- and vehicle-treated mice, and the acute metabolic response to forced treadmill running was essentially the same in both groups. Given that L-menthol treatment did not affect forced exercise, it seems likely that the reduction in wheel running was mediated via a yet to be defined central mechanism(32). Previous work from our group(6) demonstrated that L-menthol acutely increases energy expenditure in sedated mice through a TRPM8 and UCP1 dependent pathway. Our study extends these findings by showing that menthol similarly increases energy expenditure in freely moving, conscious mice.

Notably, despite the reduction in voluntary wheel running due to L-menthol treatment, the observed increase in energy expenditure, could help explain the exercise-induced improvements in indices of metabolic health. These findings might suggest that L-menthol treatment could enable a reduced volume of exercise to achieve beneficial metabolic effects. The magnitude of the training response, such as improvements in endurance capacity, metabolic health, and cardiovascular function, can vary based on exercise intensity, frequency, and duration (33). Therefore, further studies are needed to better understand the metabolic effects of L-menthol, particularly when the dose of exercise is controlled.

Finally, given that topical application of L-menthol has been shown to enhance thermogenesis (7,34), improve endurance performance (13), and reduce thermal stress (12) during physical exertion (14) in humans, further investigation into its combined effects with exercise training from the standpoint of metabolic health in humans may be warranted. In conclusion, our findings showed that while L-menthol treatment decreased voluntary wheel running distance, exercise still had a significant effect on reducing fat mass, weight gain and improving glucose tolerance. L-menthol increased energy expenditure, which may account for the observed improvements in indices of metabolic health, despite a reduction in voluntary exercise.

